# Mental fatigue impairs physical performance but not the neural drive to the muscle

**DOI:** 10.1101/2022.09.16.508212

**Authors:** Carlos Alix-Fages, Pablo Jiménez-Martínez, Daniela Souza de Oliveira, Sebastian Möck, Carlos Balsalobre-Fernández, Alessandro Del Vecchio

**Affiliations:** Applied Biomechanics and Sport Technology Research Group, Autonomous University of Madrid, Spain; Strength Training and Neuromuscular Physiology Research Center, National Strength and Conditioning Institute (ENFAF), Murcia, Spain; Research Group in Prevention and Health in Exercise and Sport (PHES), University of Valencia, Valencia, Spain; Department Artificial Intelligence in Biomedical Engineering, Friedrich-Alexander-Universität Erlangen-Nürnberg, Erlangen, Germany; Department of Exercise Science, Olympic Training and Testing Center of Hessen, Frankfurt am Main, Germany

**Author notes:** **Corresponding author** Alessandro Del Vecchio, Department Artificial Intelligence in Biomedical Engineering, Friedrich-Alexander-Universität Erlangen-Nürnberg, Henkestraße 91, 91052 Erlangen, Germany. **Funding:** No funding was received for this study.

**Keywords:** Resistance exercise, high-density surface electromyography, cognitive exertion, sport performance, muscle endurance

## Abstract

Mental fatigue (MF) does not only affect cognitive but also physical performance. This study aimed to explore the effects of MF on muscle endurance, rate of perceived exertion (RPE), and motor units’ activity. Ten healthy males participated in a randomised crossover study. The subjects attended two identical experimental sessions separated by three days with the only difference of a cognitive task (incongruent Stroop task [ST]) and a control condition (watching a documentary). Perceived MF and motivation were measured for each session at baseline and after each cognitive task. Four contractions at 20% of maximal voluntary contraction (MVIC) were performed at baseline, after each cognitive and after muscle endurance task while measuring motor units by high-density surface electromyography. Muscle endurance until failure at 50% of MVIC was measured after each cognitive task and the RPE was measured right after failure. ST significantly increased MF (p = 0.001) reduced the motivation (p = 0.008) for the subsequent physical task and also impaired physical performance (p = 0.044). However, estimates of common synaptic inputs and motor unit discharge rates as well as RPE were not affected by MF (p> 0.11). In conclusion, MF impairs muscle endurance and motivation for the physical task but not the neural drive to the muscle at any frequency bands. Although it is physiologically possible for mentally fatigued subjects to generate an optimal neuromuscular function, the altered perception and motivation seems to limit physical performance. Our results suggest that the corticospinal pathways are not affected by MF.

## INTRODUCTION

After cognitive tasks such as working (Hockey & Earle, 2006), studying (Sievertsen *et al*., 2016) or driving (Simon *et al*., 2011), humans could experience mental fatigue (MF). MF is commonly defined as a gradual and cumulative psychobiological state caused during or after demanding cognitive tasks that is characterized by physiological, behavioural, and subjective responses manifested through a lack of energy and tiredness (Okogbaa *et al*., 1994; Boksem & Tops, 2008). In this sense, during daily transport, MF is a major cause of road accidents (Lal & Craig, 2001). In the same way, pilots in United States with longer duty periods experience greater fatigue and suffer more accidents (Goode, 2003). MF can be manifested subjectively (e.g., feeling tired or lacking energy), behaviorally (e.g., impaired cognitive performance), or physiologically (e.g., alterations in brain activity) (Van Cutsem *et al*., 2017), but not all three of these areas necessarily have to be present in mentally fatigued individuals (Van Cutsem *et al*., 2017). For example, MF could not affect cognitive performance if subjects increase their effort due to motivational incentives such as money (Boksem *et al*., 2006) or avoiding more cognitive exertion tasks if they relatively increase their performance (Hopstaken *et al*., 2016). However, MF does not only affect cognitive but also physical performance (Bray *et al*., 2008).

In 1891, Angelo Mosso reported reductions in the muscle endurance of two professors measured by his ergograph after lecturing (Di Giulio *et al*., 2006). Following this initial report, other studies keep confirming the negative effects of MF on physical performance more than a century later. In this sense, a demanding cognitive task such as an incongruent Stroop task, is able to induce MF and impair time to task failure at isometric wall sit while increasing pain perception (Boat & Taylor, 2017). These negative effects of MF on physical performance have been also found for the time till failure in an isometric endurance handgrip task at 50% of maximum voluntary isometric contraction (MVIC) (Bray *et al*., 2008) and in leg extension at 20% of MVIC (Pageaux *et al*., 2013) but not for maximal force production (Bray *et al*., 2008; Pageaux *et al*., 2013). It has been proposed that the detrimental effects of MF on physical performance could be mediated by higher-than-normal levels of ratings of perceived exertion (RPE). In this sense, Marcora and colleagues found in 2009 that MF impaired the time-to-exhaustion in a cycle ergometer test and increased RPE (Marcora *et al*., 2009). Another study reported that MF impaired Yo-Yo test performance by 16%, what seemed to be mediated by the increase in RPE (Smith *et al*., 2016). This higher-than-normal RPE could be mediated by cerebral adenosine accumulation, specifically, in the anterior cingulate cortex (ACC), during cognitive demanding tasks (Martin *et al*., 2018). This accumulation of adenosine in the ACC could subsequently inhibit dopamine release, not only increasing the RPE related to exercise but also reducing the motivation to extend the effort (Martin *et al*., 2018).

MF depletes cognitive resources such as behavior regulation or decision-making influenced by the sensations perceived throughout exercise that are key for physical performance at submaximal intensities but not for maximum force production (Aitken & MacMahon, 2019). Note that previous studies found no effects of MF in maximum force production (Bray *et al*., 2008; Pageaux *et al*., 2013) or the maximal capacity of the nervous system to fully activate the muscles (Pageaux *et al*., 2013, 2015). However, when mentally fatigued individuals perform submaximal isometric handgrip tasks, the forearm muscle activity measured by electromyography (EMG) is increased (Bray *et al*., 2008; Brown & Bray, 2017). These results are in conflict with other reports showing no effects of MF on central and peripheral fatigue (Pageaux *et al*., 2015), since a higher EMG amplitude indirectly indicates higher motor unit discharge characteristics after MF. However, most of the previous studies used the global surface EMG signal, which represents a crude estimate of neural drive to the muscle (Farina *et al*., 2014*a*). With respect to the latter point, one study explored the effects of a cognitive challenge (i.e., mental math task) during a low-intensity contraction (i.e., 5% MVC), found that the increased oscillations in the common synaptic input to motor units of biceps brachii could explain the impairment in force steadiness (Pereira *et al*., 2019), although they did not explore effects on force or endurance physical performance. It is currently unknown if the effect of a cognitive challenge alters the motor unit discharge characteristics that determine a lower task performance (i.e., time to exhaustion for a given task).

To our knowledge, the effects of MF induced by an acute cognitive task on subsequent muscle endurance, motor unit activity and common inputs to motoneurons have not been explored. This is important since the oscillations carried by the motoneuron have direct link with the motor cortex (Bräcklein *et al*., 2022) and effective neural drive to the muscle (Farina *et al*., 2014*b*), and muscle performance (Del Vecchio *et al*., 2019*a*, 2019*b*).The aim of this study are was to explore the effects of MF on submaximal muscle endurance, RPE and motor unit activity. We hypothesize that MF will impair physical performance while increasing RPE and perceived mental fatigue and reducing the motivation for the subsequent physical task. Moreover, we hypothesised that these changes will be paralleled by an increase in the oscillatory activity of the motor neurons and higher discharge characteristics.

Although the MF task decreased the time to task failure, we found a similar neural drive to the muscle. Our results suggest that MF does not impar corticospinal connectivity and does not change the discharge characteristics of the motor units. The reduced time to task failure after MF is therefore determined by brain processes that result in a higher perception of effort.

## METHOD

### Participants

Ten young healthy adult males volunteered to participate in this study (age = 28.4 ± 2.7 years; body mass = 78.5 ± 12.9 kg; height = 1.81 ± 0.6 m). None of the subjects reported any physical, neurological or cardiovascular pathology that could compromise the outcomes of the present study. Subjects were instructed not to consume alcohol or any stimulating substance (e.g., caffeine) and not to perform any intense physical exercise 24 hours before each session. All subjects were informed in detail about the experimental protocol before signing the informed consent, but they were blinded with respect to the real aim of the study because this would be a potential confounding factor. The study protocol adhered to the tenets of the Declaration of Helsinki and was approved by the Institutional Review Board.

### Study design

A blinded crossover design was used to explore the effects of MF on muscle endurance performance, RPE, motor units and the common inputs to motoneurons (Figure 1). Participants attended the laboratory two times separated by 72 hours. The two experimental sessions were identical with the only difference of the cognitive task applied. The order of the two experimental sessions (STROOP TASK [ST], and CONTROL [CON]) was randomised using the Research Randomizer website (www.randomizer.org). At the beginning of each experimental session, participants reported their baseline perceived MF and motivation for the upcoming exercise task. Subsequently, electrodes were placed in tibialis anterior muscle and after the warm up, the MVIC was measured twice. At this moment, 4 trapezoidal contractions were performed at 20% of the best MVIC value for each subject while measuring the activity of the motor units by HDsEMG. After all these PRE tests, participants performed 30 minutes of the ST or CON task. After completing the high (ST) or low (CON) demanding cognitive task, their perceived MF and motivation for the upcoming exercise task were measured again. Then, the activity of the motor units during 4 trapezoidal contractions at 20% of the MVIC was measured again. At this moment, subjects performed an isometric muscle endurance task till failure at 50% of MVIC. RPE was measured right after failure. Finally, after reporting their RPE, subjects performed 4 more trapezoidal contraction at 20% of MVIC to measure their motor units activity. All sessions were held at the same time of the day for each participant and under similar environmental conditions (∼22° C and ∼60% humidity).

**Fig. 1.**
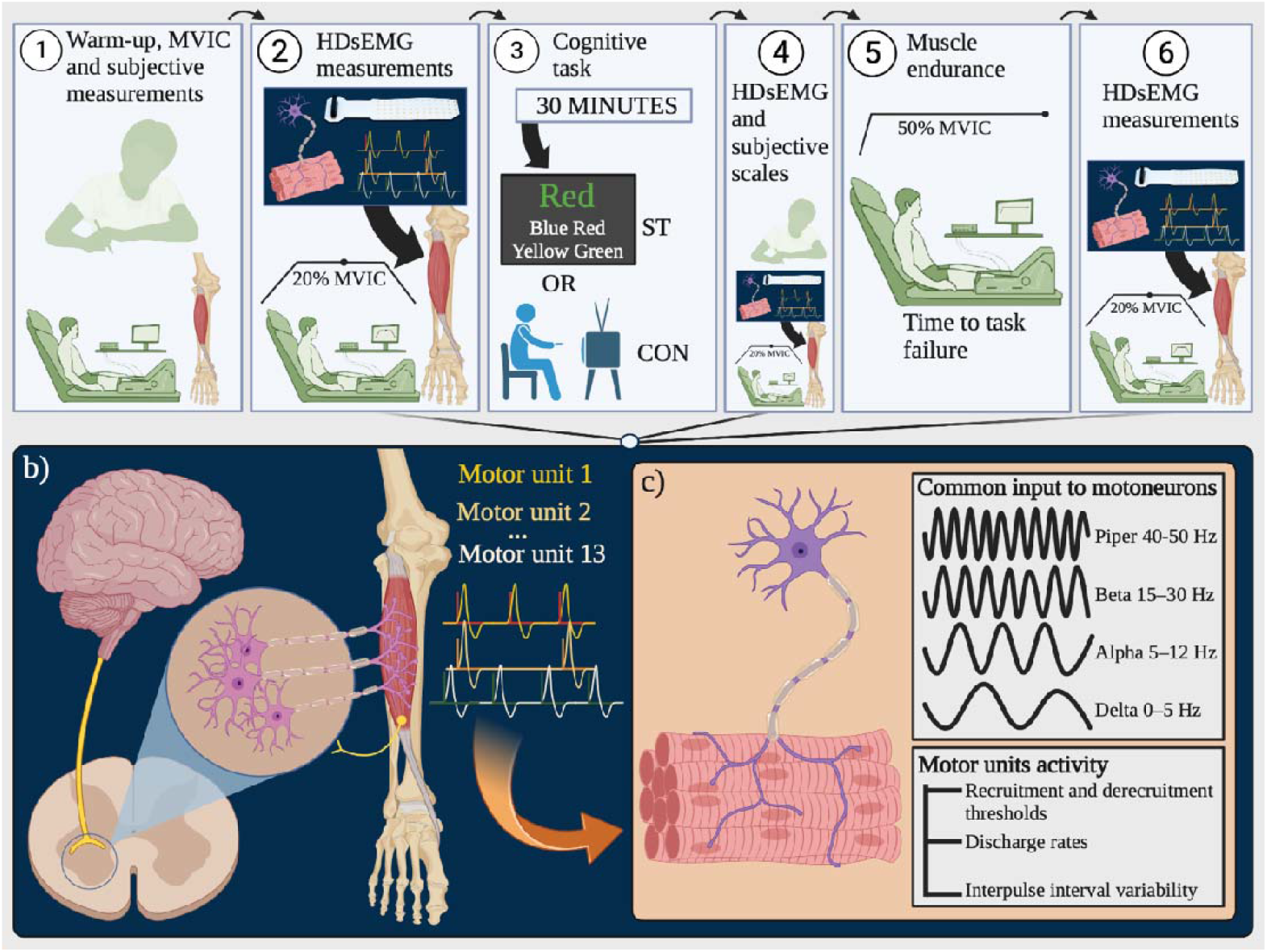
Overview of the experimental protocol (a) and methodological aspects for measured motor units of tibialis anterior (b) and outcomes obtained by HDsEMG (c).

### Procedures

#### Visual analogue scales for mental fatigue and motivation

At the beginning of each session, the subjective rate of MF was quantified by the 100mm visual analogue scale (VAS) previously adopted in the literature (Smith *et al*., 2016) to ensure that there were not differences in the levels of MF of the subjects for each session baseline. MF VAS has demonstrated to be the most practical valid method to assess MF (Smith *et al*., 2019). Note that MF is a psychophysiological state caused by prolonged periods of demanding cognitive activity and mainly characterized by subjective feelings of “tiredness” and “lack of energy” (Boksem & Tops, 2008). Besides, another 100mm VAS was used to quantify the motivation of the subjects for the upcoming exercise task (Smith *et al*., 2016). Note that theoretically, if MF is produced by the accumulation of adenosine in the ACC, it should also reduce the level of motivation due to the subsequent inhibition of dopamine release (Martin *et al*., 2018). MF and motivation levels were also reported by the participants using these VAS scales after performing the randomized intervention of each experimental session (ST or CON) to explore the effect of the high (ST) and low (CON) demanding cognitive task. The scales were anchored at one end with “none at all” and at the other end with “maximal” (Smith *et al*., 2016). No other markings were displayed on the scales. Participants were instructed to draw a mark on the continuum line at the point that best represented the feeling of their current state. The VAS score was determined by a researcher that measured the millimetres of the mark drew by each participant.

### MVIC, ramp contractions and muscle endurance

After reporting the MF and motivation baseline, participants completed a brief progressive warm-up involving a total of five isometric contractions of the ankle dorsiflexors at different intensities of self-perceived maximal voluntary force (2 × 30, 2 × 60 1 × 80% of perceived MVIC). Then, the MVIC of the ankle dorsiflexors was measured twice with 1 minute of rest between contractions. Verbal instruction to “pull as hard as possible” for 3–5 s was provided before all MVICs. Strong and loud verbal encouragement to achieve the maximum force within each contraction and to overcome the peak force of the previous MVICs was provided by an investigator. The highest force value of the two contraction was selected for the MVIC of each subject per session. MVIC was used as the reference for the submaximal intensities of the subsequent physical tasks.

Five minutes after the MVIC measurement, participants warmed up performing 3 trapezoidal (ramp) contractions at 20% of MVIC. 2 minutes after the warm-up, the real measurement of 4 trapezoidal contractions at 20% MVIC started. In this task, subjects were asked to follow a ramp showed in a monitor that represented with a red line their targeted submaximal force relative to their MVIC measured at the beginning of the session. Each of these 4 ramps started with a linear force increase during 4 seconds till reaching the 20% of their ankle dorsiflexors MVIC, which was maintained for 10 seconds (plateau). Then, subjects linearly reduced their force production following the line drop during 4 seconds. Each ramp was separated by 10 seconds rests. A real-time force recording was overlaid on the template for feedback. These 4 trapezoidal (ramp) contractions were performed by the subjects in both sessions right before the cognitive task, after the cognitive task and after the isometric muscle endurance task to failure. As explained below, these measurements were used to estimate the common input to motoneurons and motor units activity.

2 minutes after the second set of 4 trapezoidal contractions of each session, when subjects already performed the cognitive task, an isometric muscle endurance task was performed to failure. For this physical task, subjects also followed a red line depicted in the monitor that represented their relative submaximal force as explained above. In this case, subjects linearly increased their force at a rate of 5% MVIC/second till reaching their 50% MVIC. Verbal instruction to “when reaching the 50% of your maximum force, maintain it following the red line as long as you can” was provided before the task. No verbal interactions were provided during the task to not bias the results. Failure was considered when subjects could not maintain their force above 40% of MVIC. Muscle endurance performance was measured as the time to task failure in seconds.

#### Ratings of perceived exertion

We monitored RPE right immediately after subjects reached failure in the muscle endurance task of each session using a CR-10 scale (Foster *et al*., 2001). We asked subjects “How hard was the last physical task?” to quantify their perceived effort. An image of the CR-10 scale where 0 is “rest” and 10 represents “maximal” effort, was shown to the participants and they verbally reported their RPE value.

#### HDsEMG recordings

The HDsEMG signals were recorded in monopolar mode using a multichannel amplifier (Quattrocento, OT Bioelettronica, Turin, Italy), with 2048Hz sampling frequency, 16-bit analog-to-digital (A/D) converter resolution and bandpass filter 10-500Hz.

The HDsEMG signals were recorded from the tibialis anterior muscle of the dominant leg. First, the muscle perimeter was identified through palpation, the skin in this area was shaved and cleansed with 70% ethanol. After that, a grid of 64 electrodes (13 rows × 5 columns; gold-coated; diameter 1 mm; interelectrode distance 8 mm; OT Bioelettronica) was placed over the muscle belly. The grid was attached to the skin surface using a bi-adhesive foam (SpesMedica, Battipaglia, Italy) and a conductive paste (SpesMedica) to improve the contact between skin-electrodes.

As reference and ground electrodes, two strap electrodes were dampened with water. The first was positioned on the medial malleolus of the dominant leg, and the later was placed on the styloid process of the ulna was used as ground electrode.

#### Force signal recordings

The experimental setup consists of a custom-built adjustable ankle ergometer (OT Bioelettronica) secured with two straps to a massage table. The participants were seated comfortably on the table with the dominant leg extended (knee extended at 180° - neutral position), the hip was flexed at ∼120° and dominant ankle at ∼100° plantar flexion (perpendicular to the tibia). The dominant foot and ankle were secured by Velcro straps, with the foot placed on a foot plate connected in series with a load cell (CCT Transducer s.a.s, Turin, Italy), positioned perpendicular to the plantar surface of the foot. The non-dominant leg was resting on the massage table.

The analogue signal from the load cell was amplified (100x), sampled at 2048Hz and converted to digital signal with 16-bit A/D converter (Quattrocento, OT Bioelettronica). The force and HDsEMG signals were synchronized at source and acquired with the software OT BioLab (OT Bioelettronica). The same software also provided visual feedback to the participants, showing the target force templates and the force performed during the contractions on a computer monitor in front of them.

#### Force and HDsEMG analysis

The force signals were converted to newtons (N) and the offset was removed by subtracting the baseline force values. These signals were low-pass filtered with a Butterworth filter (fourth-order, zero-lag, cut-off frequency of 15 Hz). The normalized force values (%MVIC) were obtained by averaging the force values at the steady force phase of the ramp contractions at the target values.

Regarding the HDsEMG signals, first these signals were bandpass filtered at 20-500Hz Butterworth filter and then decomposed into motor unit action potentials. A convolutive blind source separation method (Holobar & Zazula, 2007) was used for automatic decomposition. After that, the detected motor unit spike trains were manually inspected, and the motor units with pulse-to-noise ratio <30dB were removed from the analysis (Del Vecchio *et al*., 2020).

Using the motor unit discharge times identified by decomposition, we computed the recruitment and derecruitment thresholds (force value in %MVIC at which the first/last motor unit action potential was discharged, respectively). The average discharge rate in pulses per second (pps) during the entire contraction, during the recruitment, steady (plateau) and derecruitment phases was also calculated for each motor unit. The average discharge rate, during recruitment and derecruitment was computed considering the average discharge times of the first/last four motor unit action potentials, respectively. For the plateau phase, the average of the first 10 action potentials was considered. The interpulse interval, which is the time interval between successive motor unit action potentials, was also computed during the steady phase of contraction.

A coherence analysis was also performed to assess the common input between the identified motor units. The coherence was calculated at the different frequencies: delta (0-5Hz), alpha (5-12Hz), beta (15-30Hz) and piper (40-50Hz). The magnitude-squared coherence was calculated using the Welch periodogram with nonoverlapping windows of 1 s. The coherence is measured from 0 to 1, with 1 indicating perfect linear relationship between two signals at that frequency and 0 indicating no relationship.

#### Control and mental fatigue experimental intervention

Two different interventions were performed by all the participants of the present study (ST and CON) in a random order. Of the two interventions, one of them was the control task while the other one was designed to induce mental fatigue performing a 30 minutes incongruent Stroop task (Smith *et al*., 2016).

30 minutes of continuous colour ST, as used in the present study, has been shown to be enough for inducing mental fatigue and impairing physical performance (Smith *et al*., 2016). For the incongruent Stroop colour word task, participants were presented with lists of colour words (i.e. “blue”, “red”, “yellow” or “green”) in the screen in which the ink colour of the letters and the colour word texted were mismatched (e.g., letters colour was green and the word text read “blue”). For each word presented, participants were required to read it but to select the ink colour of the letters and not the colour word texted. Once the participant responded, another new word appeared. The other intervention was the control task. The CON protocol consisted on watching a documentary during 30 minutes about diving in the same screen used for the ST. Subject seated in the same room and in the same place for both interventions.

### Statistical analyses

All descriptive data of the dependent variables are presented as means and standard deviations (SD). The normal distribution of the variables (Shapiro-Wilk test) and the homogeneity of the variances (Levene’s test) were confirmed (p > 0.05). The Greenhouse-Geisser correction was applied when the Mauchly’s sphericity test was significant (p < 0.05). Two-way repeated measures ANOVAs (condition [ST and CON] × time [pre and post]) were applied on the perceived MF and motivation. To ensure that there were no differences at the baseline between sessions for MVIC, pre-MF and pre-motivation, T-tests were applied. Between-condition differences in muscle endurance, the number of high-quality motor units decomposed and RPE were also investigated by T-tests. Two-way repeated measures ANOVAs (condition [ST and CON] × time [pre (baseline), pre 2 and post]) were applied on the outcomes obtained by HDsEMG. Bonferroni post-hoc corrections were used to explore significant differences. The Cohen’s d effect size (ES) with 95% confidence intervals (CI) was calculated to evaluate the magnitude of the differences using the following scale: negligible (< 0.20), small (0.20–0.49), moderate (0.50–0.79), and large (≥ 0.80) (Cohen, 1988). Statistical analyses were performed using the software package SPSS (IBM ^≤^SPSS version 25.0, Chicago, IL, USA). Statistical significance was set at p < 0.05.

## RESULTS

### Mental fatigue and motivation

Descriptive values of perceived MF and motivation are presented in Table 1. There were no differences in the pre-values of perceived MF (p = 0.757) and motivation (p = 0.416) between conditions (ST and CON), indicating similar baseline levels at the beginning of each experimental session. On the other hand, the two-way repeated measures ANOVAs revealed significant effects of condition (p = 0.001), time (p = 0.001), and condition × time interaction (p = <0.001) for the perceived MF. For the perceived motivation, significant effects were found for condition (p = 0.008) and condition × time interaction (p = 0.001) but not for time alone (p = 0.125). These results indicate that ST condition was effective to induce higher levels of MF and lower levels of motivation compared to CON.

**Table 1.**
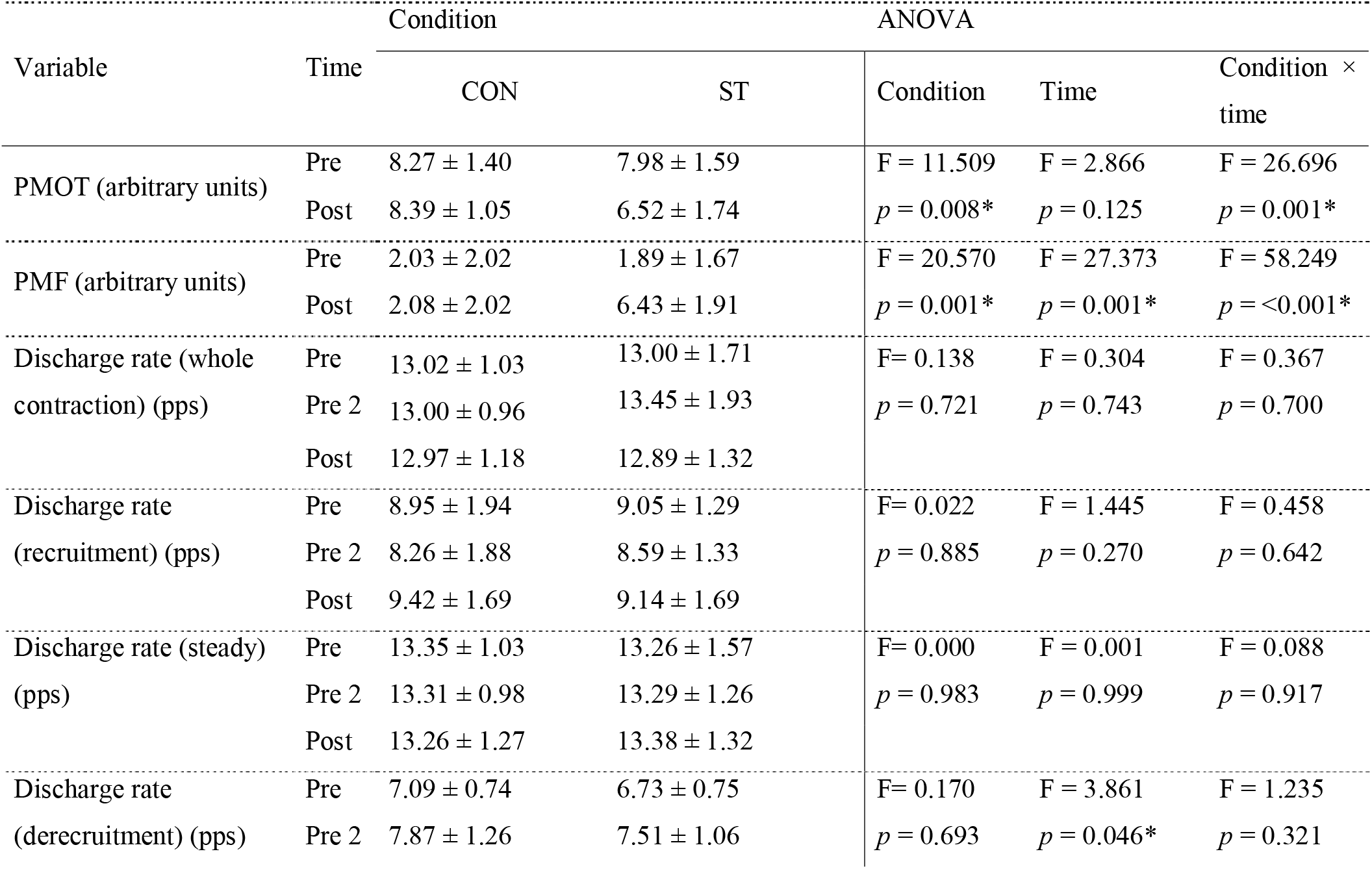
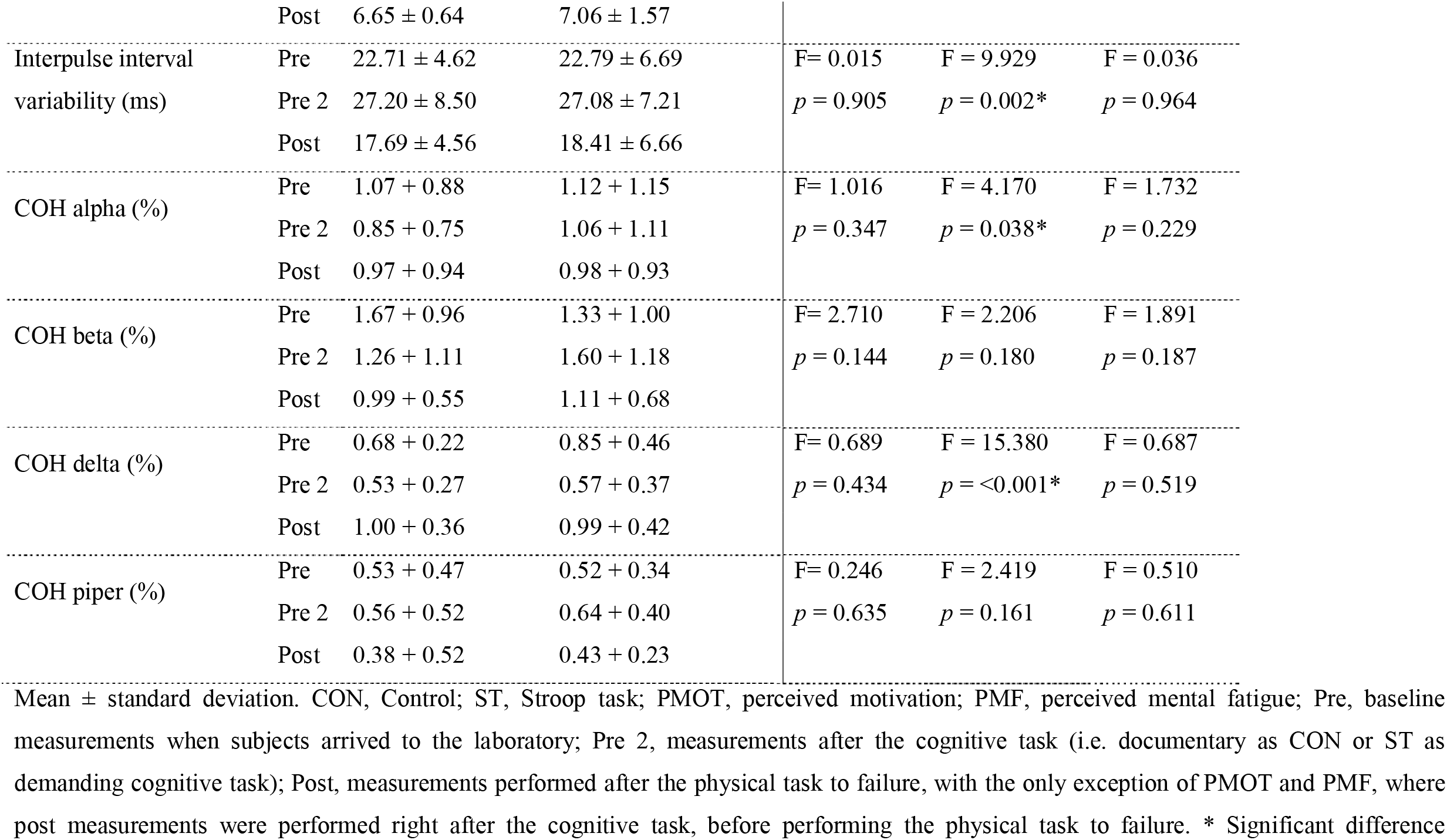
Comparison of common input to motoneurons, motor units behaviour and perceived motivation and mental fatigue between the different experimental conditions.

| Variable | Time | Condition |  | ANOVA |  |  |
| --- | --- | --- | --- | --- | --- | --- |
| | | CON | ST | Condition | Time | Condition × time |
| PMOT (arbitrary units) | Pre | $8.27 \pm 1.40$ | $7.98 \pm 1.59$ | F = 11.509 | F = 2.866 | F = 26.696 |
| | Post | $8.39 \pm 1.05$ | $6.52 \pm 1.74$ | $p = 0.008^*$ | $p = 0.125$ | $p = 0.001^*$ |
| PMF (arbitrary units) | Pre | $2.03 \pm 2.02$ | $1.89 \pm 1.67$ | F = 20.570 | F = 27.373 | F = 58.249 |
| | Post | $2.08 \pm 2.02$ | $6.43 \pm 1.91$ | $p = 0.001^*$ | $p = 0.001^*$ | $p = <0.001^*$ |
| Discharge rate (whole contraction) (pps) | Pre | $13.02 \pm 1.03$ | $13.00 \pm 1.71$ | F = 0.138 | F = 0.304 | F = 0.367 |
| | Pre 2 | $13.00 \pm 0.96$ | $13.45 \pm 1.93$ | $p = 0.721$ | $p = 0.743$ | $p = 0.700$ |
| | Post | $12.97 \pm 1.18$ | $12.89 \pm 1.32$ | | | |
| Discharge rate (recruitment) (pps) | Pre | $8.95 \pm 1.94$ | $9.05 \pm 1.29$ | F = 0.022 | F = 1.445 | F = 0.458 |
| | Pre 2 | $8.26 \pm 1.88$ | $8.59 \pm 1.33$ | $p = 0.885$ | $p = 0.270$ | $p = 0.642$ |
| | Post | $9.42 \pm 1.69$ | $9.14 \pm 1.69$ | | | |
| Discharge rate (steady) (pps) | Pre | $13.35 \pm 1.03$ | $13.26 \pm 1.57$ | F = 0.000 | F = 0.001 | F = 0.088 |
| | Pre 2 | $13.31 \pm 0.98$ | $13.29 \pm 1.26$ | $p = 0.983$ | $p = 0.999$ | $p = 0.917$ |
| | Post | $13.26 \pm 1.27$ | $13.38 \pm 1.32$ | | | |
| Discharge rate (derecruitment) (pps) | Pre | $7.09 \pm 0.74$ | $6.73 \pm 0.75$ | F = 0.170 | F = 3.861 | F = 1.235 |
| | Pre 2 | $7.87 \pm 1.26$ | $7.51 \pm 1.06$ | $p = 0.693$ | $p = 0.046^*$ | $p = 0.321$ |
|  | Post | 6.65 ± 0.64 | 7.06 ± 1.57 |  |  |  |
| Interpulse interval variability (ms) | Pre | 22.71 ± 4.62 | 22.79 ± 6.69 | F= 0.015 | F = 9.929 | F = 0.036 |
|  | Pre 2 | 27.20 ± 8.50 | 27.08 ± 7.21 | *p* = 0.905 | *p* = 0.002* | *p* = 0.964 |
|  | Post | 17.69 ± 4.56 | 18.41 ± 6.66 |  |  |  |
| COH alpha (%) | Pre | 1.07 ± 0.88 | 1.12 ± 1.15 | F= 1.016 | F = 4.170 | F = 1.732 |
|  | Pre 2 | 0.85 ± 0.75 | 1.06 ± 1.11 | *p* = 0.347 | *p* = 0.038* | *p* = 0.229 |
|  | Post | 0.97 ± 0.94 | 0.98 ± 0.93 |  |  |  |
| COH beta (%) | Pre | 1.67 ± 0.96 | 1.33 ± 1.00 | F= 2.710 | F = 2.206 | F = 1.891 |
|  | Pre 2 | 1.26 ± 1.11 | 1.60 ± 1.18 | *p* = 0.144 | *p* = 0.180 | *p* = 0.187 |
|  | Post | 0.99 ± 0.55 | 1.11 ± 0.68 |  |  |  |
| COH delta (%) | Pre | 0.68 ± 0.22 | 0.85 ± 0.46 | F= 0.689 | F = 15.380 | F = 0.687 |
|  | Pre 2 | 0.53 ± 0.27 | 0.57 ± 0.37 | *p* = 0.434 | *p* = <0.001* | *p* = 0.519 |
|  | Post | 1.00 ± 0.36 | 0.99 ± 0.42 |  |  |  |
| COH piper (%) | Pre | 0.53 ± 0.47 | 0.52 ± 0.34 | F= 0.246 | F = 2.419 | F = 0.510 |
|  | Pre 2 | 0.56 ± 0.52 | 0.64 ± 0.40 | *p* = 0.635 | *p* = 0.161 | *p* = 0.611 |
|  | Post | 0.38 ± 0.52 | 0.43 ± 0.23 |  |  |  |

Mean ± standard deviation. CON, Control; ST, Stroop task; PMOT, perceived motivation; PMF, perceived mental fatigue; Pre, baseline measurements when subjects arrived to the laboratory; Pre 2, measurements after the cognitive task (i.e. documentary as CON or ST as demanding cognitive task); Post, measurements performed after the physical task to failure, with the only exception of PMOT and PMF, where post measurements were performed right after the cognitive task, before performing the physical task to failure. \* Significant difference

### Muscle endurance performance and RPE

Significant differences were observed in the time to task failure (muscle endurance performance) between the experimental conditions (p = 0.044) with better physical performance obtained for CON (87.14 ± 44.91 seconds) condition compared to ST (75.00 ± 37.87), indicating a detrimental effect of the high demanding cognitive task on muscle endurance. However, RPE values did not reach significant differences between conditions (p = 0.111). Nonetheless, subjects tended to report higher RPE in the ST condition (5.80 ± 1.40) compared to CON (5.10 ± 1.91) with a small effect size (ES = 0.42; CI = −0.12, 0.96). A small effect size was also found for the significant differences in the time to task failure when comparing ST to CON (ES = −0.29; CI = −0.58, −0.01). The individual variability in muscle endurance and RPE are depicted in Figure 3.

### Motor unit decomposition

Subjects were excluded from motor units analyses if fewer than three high-quality motor units were decomposed in their muscle (i.e. tibialis anterior) as reported before in literature (Laine *et al*., 2015). Considering the criteria presented in 2020 by Del Vecchio et al. such as pulse-to-noise ratio to evaluate the quality of the decomposed motor units (Del Vecchio *et al*., 2020), 2 of the 10 subjects were excluded from motor units analyses due to the less than three high-quality motor units detected in their contractions. The mean ± SD number of motor units decomposed in subjects was 8.38 ± 2.56 for the control session and 8.88 ± 2.47 for the Stroop task session. T-test reported no significant differences between the number of motor units decomposed between conditions (i.e., the Stroop task session compared to control session) (p = 0.351).

### Coherence analyses for inputs to motor units

Descriptive values of COHalpha, COHbeta, COHdelta and COHpiper are presented in Table 1. The repeated measures ANOVAs revealed no significant differences for COHalpha, COHbeta, COHdelta nor COHpiper between conditions (p range = 0.144 to 0.635) or condition × time interaction (p range = 0.187 to 0.611) (Table 1). On the other hand, significant differences for time were found in COHalpha (p = 0.038) and COHdelta (p = <0.001), while no differences for time were found in COHbeta (p = 0. 0.180) and COHpiper (p = 0.161) (Table 1). However, post-hoc Bonferroni analyses did not reach any significant difference in time for COHalpha. Otherwise, post-hoc Bonferroni time analyses revealed significant higher levels of COHdelta in baseline (pre) compared to post cognitive task measurement (pre 2) (p = 0. 0.008). Besides, the highest levels of COHdelta were found at post physical task to failure measurement (post) with significant differences compared to the lower values found at post cognitive task measurement (pre 2) (p = 0. 0.009). The individual variability in common inputs to motorneurons is depicted in Figure 2.

**Fig. 2.**
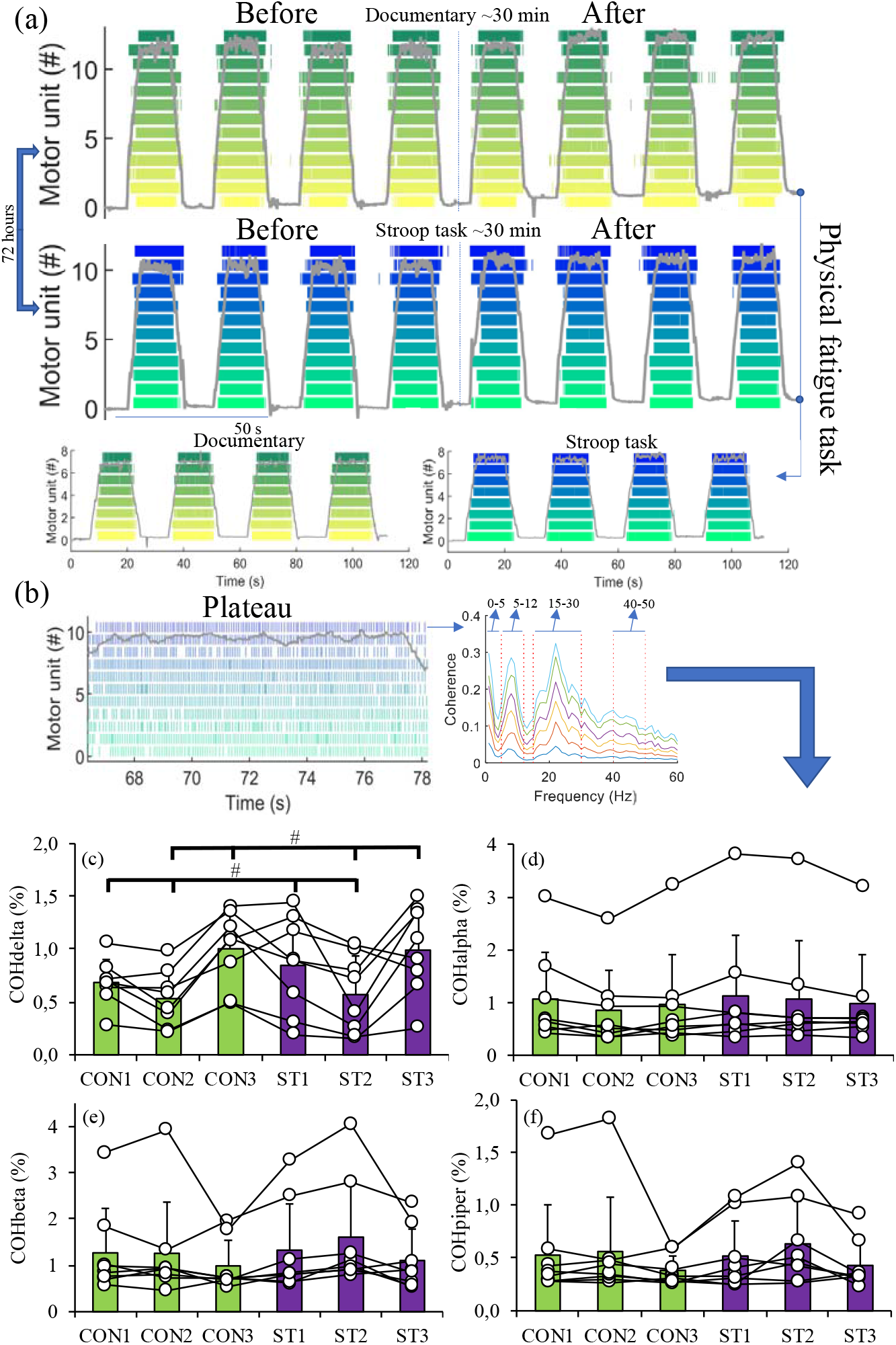
Activity of motor units before and after each cognitive task and after the subsequent physical task to failure is depicted for both sessions (ST and CON) at the beginning of the figure (a). Graphical explanation of coherence analyses during the plateau phase of the ramp contraction (b). Note that each individual line in the coherence plot indicates the coherence between an increasing number of motor unit (the smallest line indicates the coherence value between two pairs of motor units and the highest value corresponds to 14 motor units). The individual (dots), averaged across the subjects (bars) and standard deviation (line over the bars) values obtained during the ST and CON conditions for (c) COHdelta (d) COHalpha, (e) COHbeta and (f) COHpiper are represented. Three measurements were performed for each session. After CON or ST, number 1 corresponds to baseline measurements (pre), number 2 corresponds to measurements after the cognitive task (pre 2) and number 3 corresponds to measurements after the subsequent physical task to failure (post). Green bars represent CON session and blue bars represent ST session. * Significant difference between conditions (CON compared to ST). # Represents post-hoc Bonferroni significant differences between time points, present in COHdelta for pre compared to pre 2 and for post compared to pre 2.

### Motor units activity

Descriptive values of motor units activity are presented in Table 1. The repeated measures ANOVAs did not reveal significant differences between conditions (p range = 0.319 to 0.983) nor in condition × time interaction (p range = 0.165 to 0.964) for any variable (Table 1). In addition, similar results were found for recruitment and derecruitment threshold, showing not significant differences for condition (p = 0.677 and 0.967 respectively), time (p = 0.789 and 0.624 respectively) or condition × time interaction (p = 0.608 and 0.165 respectively). However, significant differences in time were found for discharge rate at derecruitment (p = 0.046) and interpulse interval variability (p = 0.002) (Table 1). Post-hoc Bonferroni analyses only showed significant differences in time for discharge rate at derecruitment after the cognitive task (pre 2) compared to the lower levels found after the subsequent physical task to failure (post) (p = 0.025). For interpulse interval variability, post-hoc Bonferroni analyses showed significantly lower levels after the physical task to failure (post) compared to both, baseline (pre) (p = 0.019) and pre 2 (after the cognitive task) (p = 0.038) measurements. These results revealed effects of physical fatigue but not mental fatigue on motor units activity. The individual variability in motor units activity outcomes is depicted in Figure 3.

**Fig. 3.**
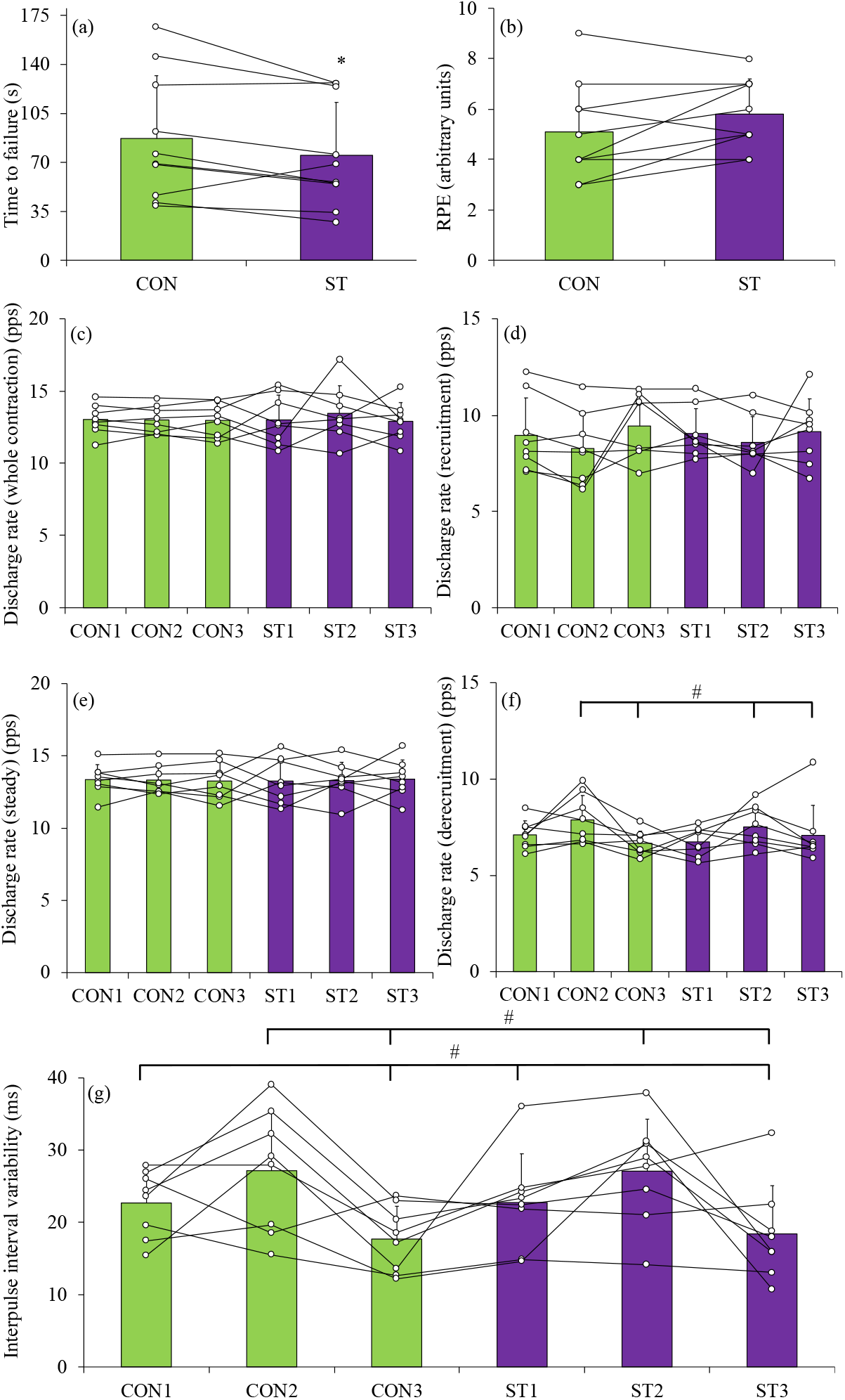
Individual (dots), averaged across the subjects (bars) and standard deviation (line over the bars) values obtained during the ST and CON conditions for (a) time to task failure (muscle endurance performance in seconds), (b) ratings of perceived exertion (RPE), (c) discharge rate (i.e. pulses per second [pps]) across the whole contraction, (d) discharge rate at recruitment, (e) discharge rate at the steady contraction, (f) discharge rate at derecruitment and (g) interpulse interval variability. For outcomes related to the activity of motor units (c-g), three measurements were performed for each session (ST and CON). After CON or ST, number 1 corresponds to baseline measurements (pre), number 2 corresponds to measurements after the cognitive task (pre 2) and number 3 corresponds to measurements after the subsequent physical task to failure (post). Green bars represent CON session and blue bars represent ST session. * Significant difference between conditions (CON compared to ST). # Represents post-hoc Bonferroni significant differences between time points, present in interpulse interval variability for post compared to both pre and pre2, and in discharge rate at derecruitment for post compared to pre 2.

## DISCUSSION

This study explored the effects of a cognitive exertion task (ST) compared to watching a documentary (CON) on the subsequent muscle endurance performance, RPE, motor units activity and their inputs from different sites such as the motor cortex or afferent feedback. The main finding of this study was that ST significantly increased MF and reduced the motivation for the subsequent physical task (i.e. time to task failure at 50% of MVIC) compared to CON but it did not affect any neurophysiological parameter measured. Although the physical fatigue task affected neurophysiological parameters such as the interpulse interval variability, discharge rate at derecruitment or COHdelta (i.e. common synaptic input in the delta band [0-5 Hz], which is related to the voluntary control of steady isometric force (Farina & Negro, 2015)), MF did not affect motor units activity nor their inputs analysed at different frequencies. These results highlight that MF impairs physical performance but not the common input to motoneurons nor neural drive to the muscle.

In line with our hypothesis, 30 minutes of continuous incongruent colour ST was significantly effective to induce MF, reduce the motivation for the subsequent physical task and finally impair muscle endurance measured as time to task failure at 50% of MVIC. These results agree with previous findings reporting that 30 minutes or less of continuous incongruent colour ST are effective to induce mental fatigue and impair physical performance (Bray *et al*., 2008; Smith *et al*., 2016; Boat & Taylor, 2017). Note that two different recent meta-analyses demonstrated that the duration of the mental effort task does not predict the magnitude of impairment, showing similar exercise performance impairments after long and short duration cognitive exertion tasks (Giboin & Wolff, 2019; Brown *et al*., 2020). On the other hand, we did not find a significant effect of ST on RPE while previously reported data did (Marcora *et al*., 2009; Smith *et al*., 2016). However, subjects tended to report higher RPE in the ST condition (5.80 ± 1.40) compared to CON (5.10 ± 1.91) with a small effect size (ES = 0.42; CI = −0.12, 0.96). In this sense, a possible explanation of this discrepancy is that while Marcora et al. 2009 and Smith et al. 2016 used physical tasks that involved the whole lower body and this may result in higher RPE values after exhaustion. However, considering that time to task failure after ST was lower than after CON, and final RPE was similar, higher-than-normal levels of RPE are present in our study after ST as the rate of increase in RPE on the physical task was greater.

Rejecting our hypothesis, none of the neurophysiological outcomes were significantly affected by ST compared to CON. In this sense, ST did not affect recruitment or derecruitment threshold, interpulse interval variability nor discharge rates at recruitment, derecruitment, steady or whole contraction.

Besides, no significant differences were found between ST and CON for any outcome obtained by coherence analyses (i.e. COHdelta, COHalpha, COHbeta and COHpiper). This means that MF did not affect motor units activity nor their common inputs analysed at different frequencies. Note that common inputs to motoneurons should be sensitive to changes in corticospinal entrainment (Ibáñez *et al*., 2022). However, the physical task significantly affected to neurophysiological parameters such as the interpulse interval variability, discharge rate at derecruitment or COHdelta. COHdelta represents the common synaptic input in the delta band (0-5 Hz), which is usually related to the voluntary control of force (Farina & Negro, 2015) and accurately reflects the force variability (Negro *et al*., 2009). In this sense, the significant increase in COHdelta after the physical task, could compensate fatigue for maintaining the recruitment of motor units and discharge rate during the steady plateau of the submaximal contractions. COHalpha represents the common synaptic input in the alpha band (5–12 Hz), which is usually associated to afferent feedback (Mehrkanoon *et al*., 2014). COHbeta represents the common synaptic input in the beta band (15-30 Hz), which is associated to descending inputs from motor cortex during isometric contractions of weak to moderate strength (Brown, 2000; Bräcklein *et al*., 2022) as performed in the present study. COHpiper represents the common synaptic input at 40-50 Hz, which is associated to descending inputs from motor cortex during strong isometric contractions or during dynamic movement (Brown, 2000). Our results differ than previous results suggesting that MF could impair physical performance by affecting corticospinal connectivity (Budini *et al*., 2014).

The hypotheses related to the effects of MF weakening the cortico-muscular coupling (i.e., synchronized activity of the motor cortex and the spinal motoneuron pool) have been partly based on data reported by Budini and colleagues in 2014 finding that mental fatigue induced by 100 minutes of a demanding cognitive task, generated a reduction in mechanically induced tremor at 8–12 Hz (Budini *et al*., 2014). Besides, another study found the increased oscillations in the common synaptic input to motor units of biceps brachii as a possible explanation of the decrease in force steadiness when performing a mental math task during a low-intensity contraction (5% MVC) (Pereira *et al*., 2019). However, several studies found no effects of MF on maximal force production or the capacity of the nervous system to fully recruit motor units (Pageaux *et al*., 2013, 2015). To our knowledge, the present study is the first exploring the effects of MF induced by prior incongruent Stroop task on subsequent muscle endurance, motor units activity and common inputs to motoneurons estimated by HDsEMG measurements, reporting no effects of MF on any of all neurophysiological parameters. Accordingly and also considering previous data reporting that MF impaired physical performance but not physiological responses such as cardiorespiratory or metabolic outcomes (Marcora *et al*., 2009), it seems that MF does not alter physiological parameters such as motor units activity, metabolic or cardiorespiratory functions.

It has been suggested that MF could impair physical performance by increasing RPE and reducing the motivation for the physical task, what could be due to the accumulation of adenosine in the ACC induced by MF (Martin *et al*., 2018). This hypothesis has been supported by data indicating that caffeine supplementation—which acts by binding to adenosine receptors—averts the negative effects of mental fatigue on exercise performance (Franco-Alvarenga *et al*., 2019; Grgic, 2021). Based on these findings and our data indicating that MF significantly reduces the motivation for the subsequent physical task and tends to increase RPE, we suggest that MF impairs physical performance by altering prefrontal cortex circuitry but not the corticospinal connectivity. In this sense, although it is physiologically possible for the motor units to generate an optimal neuromuscular function, the altered perceptions and motivation due to changes at prefrontal cortex levels triggers into an unconscious decision of reduce the magnitude of the neuromuscular effort or stop the physical task. Note that he ACC is linked to areas of the prefrontal cortex such as the dorsolateral prefrontal cortex (DLPFC) (Tang *et al*., 2019), which is also affected by mental fatigue (Müller & Apps, 2019). The DLPFC plays a key role in the regulation and integration of perceptual responses, the capacity to tolerate high levels of physical exertion, effort-based decision-making, self-regulatory processes during physical effort and possibly in the determination of exercise termination (Robertson *et al*., 2016; Wolff *et al*., 2018). Note that these cognitive resources depleted by MF are key for physical performance at submaximal intensities (Aitken & MacMahon, 2019). In this sense, non-invasive brain stimulation over the DLPFC has improved strength endurance performance while reducing RPE (Alix-Fages *et al*., 2020). On the other hand, in line with the absence of MF effects on power tasks, non-invasive brain stimulation over the DLPFC does not significantly affect power tasks performance such as CMJ (Romero-Arenas *et al*., 2019), MVC (Alix-Fages *et al*., 2019) or sprint (Alix-Fages *et al*., 2021, 2022).

The present study is not free of limitations. Although, every subject participated in both experimental conditions and as controlled by pre and post measurements of each session, larger sample size may benefit the generalizability of our preliminary report. Besides, our results could not be extrapolated to highly trained athletes. In conclusion, MF impairs muscle endurance performance and the motivation for the physical task but not motor units activity nor common inputs to motoneurons. Although it is physiologically possible for mentally fatigued subjects to generate an optimal neuromuscular function, the altered perceptions and motivation seems to limit physical performance.

## Acknowledgement

We would like to thank all the volunteers that participated in the study.

## Abbreviations

HDsEMG: high-density surface electromyography
MF: mental fatigue
MVIC: maximum voluntary isometric contraction
RPE: ratings of perceived exertion
ACC: anterior cingulate cortex
EMG: electromyography
ST: stroop task
VAS: visual analogue scale
PPS: pulse per second

## Notes

**Conflict of Interest:** Authors declare that they have no conflict of interest.

### Competing Interest Statement

The authors have declared no competing interest.

## REFERENCES

Aitken B & MacMahon C (2019). Shared Demands Between Cognitive and Physical Tasks May Drive Negative Effects of Fatigue: A Focused Review. Front Sport Act Living 1, 45.

Alix-Fages C, García-Ramos A, Calderón-Nadal G, Colomer-Poveda D, Romero-Arenas S, Fernández-del-Olmo M & Márquez G (2020). Anodal transcranial direct current stimulation enhances strength training volume but not the force–velocity profile. Eur J Appl Physiol 120, 1881–1891.

Alix-Fages C, Garcia-Ramos A, Romero-Arenas S, Nadal GC, Jerez-Martínez A, Colomer-Poveda D & Márquez G (2021). Transcranial Direct Current Stimulation Does Not Affect Sprint Performance or the Horizontal Force-Velocity Profile. Res Q Exerc SportOnline ahead of print.

Alix-Fages C, Romero-Arenas S, Calderón-Nadal G, Jerez-Martínez A, Pareja-Blanco F, Colomer-Poveda D, Márquez G & Garcia-Ramos A (2022). Transcranial direct current stimulation and repeated sprint ability: No effect on sprint performance or ratings of perceived exertion. Eur J Sport Sci 22, 569–578.

Alix-Fages, Romero-Arenas, Castro-Alonso, Colomer-Poveda, Río-Rodriguez, Jerez-Martínez, Fernandez-del-Olmo & Márquez (2019). Short-Term Effects of Anodal Transcranial Direct Current Stimulation on Endurance and Maximal Force Production. A Systematic Review and Meta-Analysis. J Clin Med 8, 536.

Boat R & Taylor IM (2017). Prior self-control exertion and perceptions of pain during a physically demanding task. Psychol Sport Exerc 33, 1–6.

Boksem MAS, Meijman TF & Lorist MM (2006). Mental fatigue, motivation and action monitoring. Biol Psychol 72, 123–132.

Boksem MAS & Tops M (2008). Mental fatigue: Costs and benefits. Brain Res Rev 59, 125–139.

Bräcklein M, Barsakcioglu DY, Del Vecchio A, Ibáñez J & Farina D (2022). Reading and Modulating Cortical β Bursts from Motor Unit Spiking Activity. J Neurosci 42, 3611–3621.

Bray SR, Martin Ginis KA, Hicks AL & Woodgate J (2008). Effects of self-regulatory strength depletion on muscular performance and EMG activation. Psychophysiology 45, 337–343.

Brown DMY & Bray SR (2017). Effects of Mental Fatigue on Physical Endurance Performance and Muscle Activation Are Attenuated by Monetary Incentives. J Sport Exerc Psychol 39, 385–396.

Brown DMY, Graham JD, Innes KI, Harris S, Flemington A & Bray SR (2020). Effects of Prior Cognitive Exertion on Physical Performance: A Systematic Review and Meta-analysis. Sport Med 50, 497–529.

Brown P (2000). Cortical drives to human muscle: The Piper and related rhythms. Prog Neurobiol 60, 97–108.

Budini F, Lowery M, Durbaba R & De Vito G (2014). Effect of mental fatigue on induced tremor in human knee extensors. J Electromyogr Kinesiol 24, 412–8.

Cohen J (1988). Statistical power analysis for the behavioral sciences, 2nd edn. Lawrence Erlbaum Associates, Hillsdale, N.J.

Van Cutsem J, Marcora S, De Pauw K, Bailey S, Meeusen R & Roelands B (2017). The Effects of Mental Fatigue on Physical Performance: A Systematic Review. Sport Med 47, 1569–1588.

Farina D, Merletti R & Enoka RM (2014a). The extraction of neural strategies from the surface EMG: an update. J Appl Physiol 117, 1215–1230.

Farina D & Negro F (2015). Common Synaptic Input to Motor Neurons, Motor Unit Synchronization, and Force Control.

Farina D, Negro F & Dideriksen JL (2014b). The effective neural drive to muscles is the common synaptic input to motor neurons. J Physiol 592, 3427–3441.

Foster C, Florhaug JA, Franklin J, Gottschall L, Hrovatin LA, Parker S, Doleshal P & Dodge C (2001). A New Approach to Monitoring Exercise Training. J Strength Cond Res 15, 109–15.

Franco-Alvarenga PE, Brietzke C, Canestri R, Goethel MF, Hettinga F, Santos TM & Pires FO (2019). Caffeine improved cycling trial performance in mentally fatigued cyclists, regardless of alterations in prefrontal cortex activation. Physiol Behav 204, 41–48.

Giboin LS & Wolff W (2019). The effect of ego depletion or mental fatigue on subsequent physical endurance performance: A meta-analysis. Perform Enhanc Heal 1, 100150.

Di Giulio C, Daniele F & Tipton CM (2006). Angelo Mosso and muscular fatigue: 116 Years after the first congress of physiologists: IUPS commemoration. Am J Physiol - Adv Physiol Educ 30, 51–57.

Goode JH (2003). Are pilots at risk of accidents due to fatigue? J Safety Res 34, 309–313.

Grgic J (2021). Effects of Caffeine on Resistance Exercise: A Review of Recent Research. Sport Med Published online.

Hockey GRJ & Earle F (2006). Control over the scheduling of simulated office work reduces the impact of workload on mental fatigue and task performance. J Exp Psychol Appl 12, 50–65.

Holobar A & Zazula D (2007). Multichannel blind source separation using convolution Kernel compensation. IEEE Trans Signal Process 55, 4487–4496.

Hopstaken JF, van der Linden D, Bakker AB, Kompier MAJ & Leung YK (2016). Shifts in attention during mental fatigue: Evidence from subjective, behavioral, physiological, and eye-tracking data. J Exp Psychol Hum Percept Perform 42, 878–889.

Ibáñez J, Zicher B, Brown KE, Rocchi L, Casolo A, Del Vecchio A, Spampinato D, Vollette C, Rothwell JC, Baker SN & Farina D (2022). Standard intensities of transcranial alternating current stimulation over the motor cortex do not entrain corticospinal inputs to motor neurons. J PhysiolOnline ahead of print.

Laine CM, Martinez-Valdes E, Falla D, Mayer F & Farina D (2015). Motor neuron pools of synergistic thigh muscles share most of their synaptic input. J Neurosci 35, 12207–12216.

Lal SKL & Craig A (2001). A critical review of the psychophysiology of driver fatigue. Biol Psychol 55, 173–194.

Marcora SM, Staiano W & Manning V (2009). Mental fatigue impairs physical performance in humans. J Appl Physiol 106, 857–864.

Martin K, Meeusen R, Thompson KG, Keegan R & Rattray B (2018). Mental Fatigue Impairs Endurance Performance: A Physiological Explanation. Sport Med 48, 2041–2051.

Mehrkanoon S, Breakspear M & Boonstra TW (2014). The reorganization of corticomuscular coherence during a transition between sensorimotor states. Neuroimage 100, 692–702.

Müller T & Apps MAJ (2019). Motivational fatigue: A neurocognitive framework for the impact of effortful exertion on subsequent motivation. Neuropsychologia 123, 144–151.

Negro F, Holobar A & Farina D (2009). Fluctuations in isometric muscle force can be described by one linear projection of low-frequency components of motor unit discharge rates. J Physiol 587, 5925–5938.

Okogbaa OG, Shell RL & Filipusic D (1994). On the investigation of the neurophysiological correlates of knowledge worker mental fatigue using the EEG signal. Appl Ergon 25, 355–365.

Pageaux B, Marcora SM & Lepers R (2013). Prolonged mental exertion does not alter neuromuscular function of the knee extensors. Med Sci Sports Exerc 45, 2254–64.

Pageaux B, Marcora SM, Rozand V & Lepers R (2015). Mental fatigue induced by prolonged self-regulation does not exacerbate central fatigue during subsequent whole-body endurance exercise. Front Hum Neurosci 9, 67.

Pereira HM, Schlinder-DeLap B, Keenan KG, Negro F, Farina D, Hyngstrom AS, Nielson KA & Hunter SK (2019). Oscillations in neural drive and age-related reductions in force steadiness with a cognitive challenge. J Appl Physiol 126, 1056–1065.

Robertson C V., Marino FE, Meeusen R, Pires FO, Pinheiro FA, Lutz K, Cheung SS, Perrey S, Radel R, Brisswalter J, Rauch HGL, Micklewright D, Beedie C & Hettinga F (2016). A role for the prefrontal cortex in exercise tolerance and termination. J Appl Physiol 120, 464–466.

Romero-Arenas S, Calderón-Nadal G, Alix-Fages C, Jerez-Martínez A, Colomer-Poveda D & Márquez G (2019). Transcranial Direct Current Stimulation Does Not Improve Countermovement Jump Performance in Young Healthy Men. J Strength Cond ResOnline ahead of print.

Sievertsen HH, Gino F & Piovesan M (2016). Cognitive fatigue influences students’ performance on standardized tests. Proc Natl Acad Sci U S A 113, 2621–2624.

Simon M, Schmidt EA, Kincses WE, Fritzsche M, Bruns A, Aufmuth C, Bogdan M, Rosenstiel W & Schrauf M (2011). Eeg alpha spindle measures as indicators of driver fatigue under real traffic conditions. Clin Neurophysiol 122, 1168–1178.

Smith MR, Chai R, Nguyen HT, Marcora SM & Coutts AJ (2019). Comparing the Effects of Three Cognitive Tasks on Indicators of Mental Fatigue. J Psychol 153, 759–783.

Smith MR, Coutts AJ, Merlini M, Deprez D, Lenoir M & Marcora SM (2016). Mental fatigue impairs soccer-specific physical and technical performance. Med Sci Sports Exerc 48, 267–276.

Tang W, Jbabdi S, Zhu Z, Cottaar M, Grisot G, Lehman JF, Yendiki A & Haber SN (2019). A connectional hub in the rostral anterior cingulate cortex links areas of emotion and cognitive control. Elife 8, e43761.

Del Vecchio A, Casolo A, Negro F, Scorcelletti M, Bazzucchi I, Enoka R, Felici F & Farina D (2019a). The increase in muscle force after 4 weeks of strength training is mediated by adaptations in motor unit recruitment and rate coding. J Physiol 597, 1873–1887.

Del Vecchio A, Holobar A, Falla D, Felici F, Enoka RM & Farina D (2020). Tutorial: Analysis of motor unit discharge characteristics from high-density surface EMG signals. J Electromyogr Kinesiol 53, 102426.

Del Vecchio A, Negro F, Holobar A, Casolo A, Folland JP, Felici F & Farina D (2019b). You are as fast as your motor neurons: speed of recruitment and maximal discharge of motor neurons determine the maximal rate of force development in humans. J Physiol 597, 2445–2456.

Wolff W, Bieleke M, Hirsch A, Wienbruch C, Gollwitzer PM & Schüler J (2018). Increase in prefrontal cortex oxygenation during static muscular endurance performance is modulated by self-regulation strategies. Sci Rep 8, 1–10.

